# A benchmarking of deep neural network models for cancer subtyping using single point mutations

**DOI:** 10.1101/2022.07.24.501264

**Authors:** Pouria Parhami, Mansoor Fateh, Mohsen Rezvani, Hamid Alinejad Rokny

## Abstract

It is now well-known that genetic mutations contribute to development of tumors, in which at least 15% of cancer patients experience a causative genetic abnormality including *De Novo* somatic point mutations. This highlights the importance of identifying responsible mutations and the associated biomarkers (e.g., genes) for early detection in high-risk cancer patients. The next-generation sequencing technologies have provided an excellent opportunity for researchers to study associations between *De Novo* somatic mutations and cancer progression by identifying cancer subtypes and subtype-specific biomarkers. Simple linear classification models have been used for somatic point mutation-based cancer classification (SMCC); however, because of cancer genetic heterogeneity (ranging from 50% to 80%), high data sparsity, and the small number of cancer samples, the simple linear classifiers resulted in poor cancer subtypes classification. In this study, we have evaluated three advanced deep neural network-based classifiers to find and optimized the best model for cancer subtyping. To address the above-mentioned complexity, we have used pre-processing clustered gene filtering (CGF) and indexed sparsity reduction (ISR), regularization methods, a Global-Max-Pooling layer, and an embedding layer. We have evaluated and optimized the three deep learning models CNN, LSTM, and a hybrid model of CNN+LSTM on publicly available TCGA-DeepGene dataset, a re-formulated subset of The Cancer Genome Atlas (TCGA) dataset and tested the performance measurement of these models is 10-fold-cross-validation accuracy. Evaluating all the three models using a same criterion on the test dataset revealed that the CNN, LSTM, and CNN+LSTM have 66.45% accuracy, 40.89% accuracy, and 41.20% accuracy in somatic point mutation-based cancer classification. Based on our results, we propose the CNN model for further experiments on cancer subtyping based on DNA mutations.

## 1. Introduction

Cancer is one of the most dangerous diseases among humans, with a mortality rate of nearly 10 million deaths in 2020[2]. Cancer can disrupt the process of birth and death of cells in the body, resulting in many cells of the body that causes tumor tissue/cell. Therefore, rapid diagnosis of cancer can effectively improve patient health.

Machine Learning (ML) techniques displayed promising ability in prediction and classification tasks [3–9], including disease prediction, virus genome analysis [10–13], and cancer subtyping. For instance, Huang et al. colorectal cancer [14], Aaroe et al. breast cancer [15], and Winnepenninckx et al. melanoma cancer [16] try to early detection cancer. Rezaei et al also provided a web-based toolset to identify breast cacner biomakrers [17].

In recent decades, easy and fast access to DNA sequences has made SMCC as one of the notable methods for cancer type identification. A DNA sequence contains many genes, all of which are not useful in diagnosing cancer and do not give us specific information. Researchers have recently used DNA information for both cancer subtyping and cancer biomarker discovery [1, 16, 18–23]. However, using machine learning methods for assimilation with SMCC bring some new challenges that have not yet been solved. One of these challenges is finding genes that are effective in identifying cancer. To solve this problem, we must find a way to identify influential and essential genes and put them in a separate group. In this regard, works have been done such as Cho et al. [24] used the mean and standard deviation of the distances from each sample to the class center as criteria for classification. Cai et al. [25] have used clustering gene selection to identify driver genes. I.e., those genes that have significant roles in cancer developments. Nevertheless, these methods do not work well on somatic point mutations that contain binary data [1]. Binary data contain values as one and zero. One represents a mutation in the gene, and zero represents no mutation in the gene.

The second problem is to identify the best genes as the features to classify cancer patients. As we mentioned before, the values in our data are one and zero. Because of the large number of zeros in our dataset, the data is sparse and difficult to categorize. The third problem is that genes have different effects on each type of cancer. Also, they have complex relationships with each other. For these reasons, simple linear classifiers like SVM [26], which are doing classification tasks in machine learning, do not work well on the SMCC problems.

To address the above-mentioned challenges, Yuan et al. [1] have invented two unique pre-processing methods called clustered gene filtering (CGF) and indexed sparsity reduction (ISR) to select effective genes and reduce sparsity in the data. They also use a deep neural network (DNN) classifier to solve a classification problem. This novel SMCC method is called DeepGene [1]. They also created a dataset named TCGA-DeepGene and used it in their experiments. [1] used 10-fold cross-validation accuracy on the training set for parameter optimization, and for comparing their work with others, they adopted test accuracy as the evaluation metric. DeepGene has achieved 65.5% test accuracy on the test dataset [1].

We perform the pre-proccesing steps based on the information provided by Yuan et al. [1]. We examine three deep learning models (CNN, LSTM, and CNN + LSTM) for identifying cancer types.

We also use Yuan et al. [1] evaluation metrics to compare our results with the results achieved in their study. We develop the models in four steps. The first and second steps consist of tuning various hyper-parameters with the help of the Keras-tuner library. We monitor metrics such as training accuracy, validation accuracy, training loss, and validation loss to evaluate the results of hyper-parameter tuning. The third and fourth steps consist of training and testing our models using the 10-fold cross-validation method.

We use regularization methods to deal with the overfitting problem in our models. The fully-connected layer is prone to overfitting problems in the CNN model. For this reason, we use global pooling instead of multi fully-connected layers. Also, to improve the performance of the models, we use the embedding layer to reduce data sparsity and place similar features in a group. In general, the challenges we face and intend to are:

– High-level data are sparse.
– Overfitting problem on our models.
– Selection of genes that have a more significant impact on classification work.

In summary, we make the following contributions:

– We examine three models of deep learning
– We introduce a CNN model to cancer classification based on somatic point mutation.
– We set a range of parameters to build our model using Keras-tuner.
– We use Global Pooling to replace multiple fully-connection layers and prevent overfitting.
– We used the embedding layer to eliminate data sparsity (a large number of zeros) and place similar features next to each other.

The rest of this paper has been organized as bellow:

Section 3 describes the basic concepts of deep neural networks. We present the related work Section. 2. Section 4 describes our approach and other properties, such as pre-processing and embedding layers. Section 5 describes our implementation and experimental results. Section 6 presents the discussion. Finally, we concluded the paper concluded in Sect. 7.

## 2. Related Works

This section will examine machine learning applications and deep learning in different cancer types/subtypes prediction and tumor diagnosis.

Many research has been done to predict the types/subtypes of cancer using machine learning methods [18–21, 27–31]. In gene expression analysis, the number of genes is usually much higher than the samples. For this reason, gene extraction methods try to increase the performance of classifiers by extracting less but more effective genes. The two main models of gene extraction are filtering and clustering models. By combining these two, Chow et al. [27] proposed a new algorithm called Double-thresholding Extraction of Feature Gene (DEFG). P. Browne et al. [29] introduced a Gaussian and uniform distributions mixture. This mixture model using in clustering and classification tasks. Sujitha et al. [30] used apache spark and a combination of binary classifier (SVM with radial basis function) and multiclass classifier (winner-take-all with SVM) to diagnose lung cancer in the early stage. They classify the data as malignant and benign and determine the extent of malignancy. Sujitha et al. (Sujitha & Seenivasagam, 2021) used sputum cells datasets and achieved 86% accuracy.

Many researchers have used the CNN model to classify cancer in their research [32–42]. Coudray et al. [35] used Incept v3 [43], one of CNN’s architectures, to diagnose two types of lung cancer Adenocarcinoma (LUAD) and squamous cell carcinoma (LUSC). EMS-Net [37] is pre-trained DenseNet-161, ResNet-152, and ResNet-101 try to classify hematoxylin-eosin stained breast histopathological microscopy images. Khan et al. [33] presented a new framework based on deep learning to diagnose and classify breast cancer in cytology images. They increased their dataset data to prevent overfitting [44, 45] and used transfer learning in their CNN architecture [46, 47]. Their model is the combination of GoogleNet [48], VGGNet [49], and ResNet [50].

Jia et al. [51] proposed CNN-SVM and a novel feature extraction method named Gabor transformation and Gray-Level Co-occurrence Matrix (GLCM) for cervical cell classification. GLCM is a statistical analysis method. It can show information of image gray surface about direction, adjacent interval, and variation range. Gabor transformation is very sensitive to the local space. Gabor can record the spatial frequency of different directions in the local area and some structural features. They use the Herval dataset and achieve 99.3% accuracy for 2-class detection.

Liang et al. [52] presented a combination of CNN (Xception [53]) and LSTM to diagnose and classify blood cells. They used the BCCD database and achieved 90.79% accuracy. This paper showed the possible combination of CNN with other methods on images, which allows us to apply such a combination on somatic point mutation data to identify cancer types. Yuan et al. [1] proposed two novel pre-processing and fully-connected neural networks for cancer classification based on somatic point mutations.

The work we examined in this section was done on images, had large datasets, or generated data to expand their datasets. However, our database data is binary (zeros and ones), the data are scattered, the number of genes is high, and the number of samples in the database is small. Genes also have complex relationships with each other that make classification difficult. These differences make the methods discussed in this section not work well on our database.

To solve these problems, based on the information reported by DeepGene, we will use the database created by them. We will compare the results obtained in this paper with their results reasonably. They also introduced two pre-processors to solve data scattering and selecting genes influencing the classification process, which we will use. We test three deep learning models called CNN, LSTM and combine these two models. To reduce the effect of data scattering, we used an embedding layer in our models. Also, due to the lack of data, we try to identify the effective genes more accurately to reduce the number of genes. To do this, we used the CNN model. Moreover, we test the performance of this model when combined with LSTM.

## 3. Backgrounds

In this section, we will briefly explain the deep neural network.

### 3.1 Deep Learning

Due to the deep hierarchical structure of the human speech system and the production and study of artificial networks (ANNs), the concept of deep learning was introduced in the late twentieth century. The remarkable breakthrough in deep learning came about when Hinton introduced a new deep learning architecture called the deep belief network (DBN) in the Year 2006 [54]. Unlike traditional approaches to machine learning and artificial intelligence, deep learning technologies have received significant advances in speech recognition applications, natural language processing, computer vision, image analysis, health care, and information retrieval [55].

### 3.2 CNN

CNN’s were the first successful deep learning architecture because of their hierarchical structure. CNN reduces the number of network parameters by empowering specific relationships and improves performance using standard back-propagation algorithms. Another feature of CNN is that it requires minimal pre-processing. CNN’s have performed well in handwriting recognition, face detection, behavior recognition, speech recognition, recommender systems, image classification, and NLP [56]. Also, CNN structures and the combination of CNN with other methods have been used successfully in tasks such as segmentation [57] or classification of medical images [58] such as breast cancer diagnosis [59, 60] or epithelial tissue detection in prostate cancer.

#### 3.2.1 CNN structure and formula

CNN has three main layers, convolutional layer (CONV), pooling layer (POOL), and fully connected layer (FC). The convolutional [61] layer and pooling layer are at the beginning of the model, and one or more fully connected layers are at the end of the model. A typical CNN usually uses a feed-forward design, where each layer uses the previous layer’s output [62, 63].

#### 3.2.2 CNN Convolutional layer

The convolutional layer is the layer that has the most use in CNN, and most of the calculations are the responsibility of this layer. The primary role of the convolutional layer is to extract features from the image or feature maps created by previous layers. It then creates the feature map output. One or more filters/kernels with *s* stride size are applied to the input to create a feature map output. Input is a color image with size(*h_in_* * *w_in_* * *d*), which is the image’s height, width, and depth or color Channel, respectively. Filter size or kernel size is(*f_h_* * *f_w_* * *d*). *f_h_* is filter’s heigh, *f_w_* is the filter’s width, and *d* is depth or channel. The output shape of the convolution layer calculate by formula (1) [62, 63]:

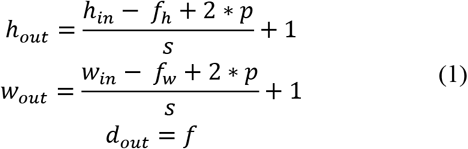

In this formula, *f* = *f_h_* = *f_w_*, and *p* is padding. Activation function could be sigmoid tanh or Relu[62, 63].

#### 3.2.3 CNN (Pooling layer)

After one or more convolutional layers, usually pooling layer is used. This layer reduces the dimensions of the feature map output with a summation of the surrounding outputs, preventing much processing, and essential features remain. The two most commonly used pooling layer methods used in CNN models are Max pooling and Average pooling. Average pooling averages the value of an area of the feature map, and max pooling selects the maximum value in an area of the feature map pooling layers shown in Figure 1.

**Figure 1.**
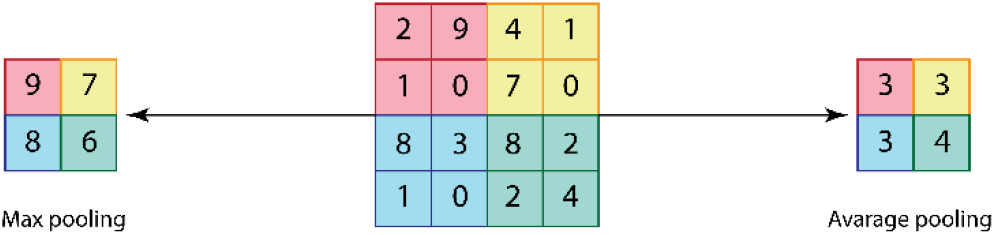
the results of applying Max pooling and Average pooling to the input matrix. Suppose the middle matrix is the input matrix, the left matrix is the result of applying Max pooling to the middle matrix, and the right matrix is the result

**Figure 2.**
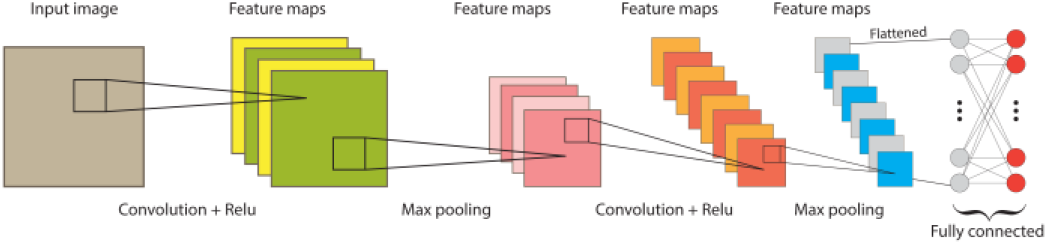
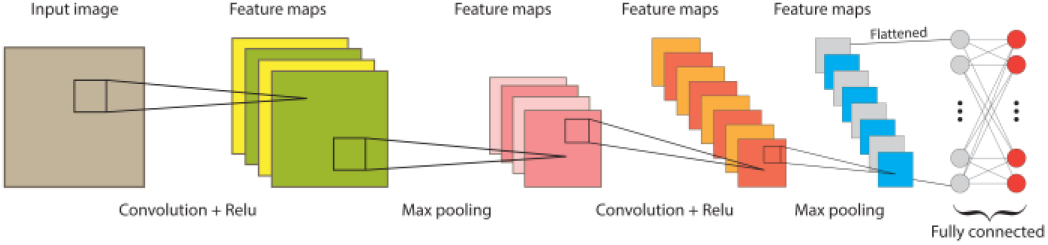
Complete structure of the standard CNN model, Parts convolution + Relu and Max pooling can be repeated several times as needed. The fully connected layer can also have the appropriate depth to the desired problem.

The pooling layer output shape calculates by formula (2). The input of the pooling layer is the output of the convolution layer, which is (*h_in_* * *w_in_* * *d_in_*) *h_in_* height, *w_in_* width, and *d_in_* is depth or channel of input data. *h_out_*, *w_out_*, *d_out_* are the output’s height, width, and channel or depth, respectively.

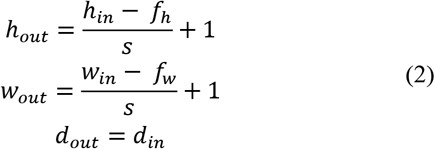

#### 3.2.4 CNN (Fully connected layer)

A fully connected layer usually appears at the end of the model and constructs the final classification. Feature maps must be converted to vectors and then given to FC.

Figure shows the complete structure of CNN [62, 63]

### 3.3 LSTM

Long short-term memory(LSTM) can learn to bridge minimal time lags over 1000 discrete time steps by enforcing constant error flow-through “constant error carrousels”(CECs) within special units called cells. Multiplicative gate units learn to open and close access to the cells. The LSTM learning algorithm is local in space and time; its computational complexity per time step and weight are O (1). LSTM solves complex long-time lag tasks that previous RNN algorithms could not solve [64].

#### 3.3.1 LSTM structure and formula

Cell LSTM has three gates called input, forget, and output. These gates have the task of selecting the signal that should go to the next node. Figure 3 shows an image of an LSTM cell, and formula (1) shows the equation of one cell in a one-time step [65, 66]. In this formula *x_t_* is the input vector, *f_t_* is the forget gate, *o_t_* is output gate, *i_t_* is input gate *h_t_* is hidden state vector, *c_t_* is cell state, 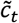 represents a candidate for cell state at timestamp(t), *σ_g_* is sigmoid activation function, *σ_c_* hyperbolic tangent activation function, and *b* represent biase. W and U are the weights of input and recurrent connections, respectively. *h*_(*t*–1)_ and *c*_(*t*–1)_ are the inputs from the previous timestep. In a cell, *x_t_*, *h_t_*, and *c_t_* are inputs, and the first step is to throw away some information from *c_t_*. This decision is made by *f_t_*. The second step is to store some new information in *c_t_*. This decision is made by two-step. First, *i_t_* selects the values for update, the second 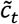 creates a vector of new candidates to state. *c_t_* update the cell state, by multiplying the *f_t_* and previuse cell state *C_(t-1)_*. Then add to 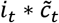. Finally, LSTM cell uses *o_t_* and *h_t_* to decide what parts of the cell state will output and create it. LSTM cell sends the *c_t_* and *h_t_* for the next step LSTM.

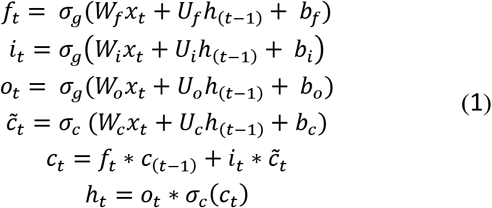

**Figure 3.**
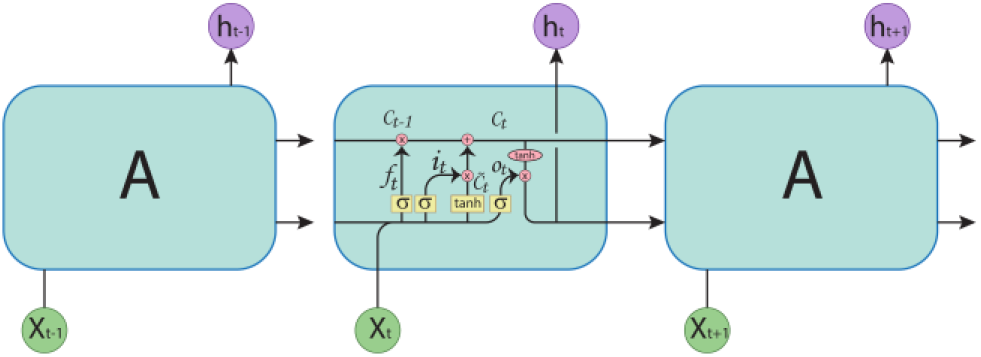
three LSTM cells and view inside one of them that can see LSTM gates.

## 4. Method and Materials

In this section, we first describe our solution then explain the details of the components.

### 4. 1 Solution overview

DeepGene [1] has used ISR and CGF pre-processing to address the two challenges of sparsity data and finding genes involved in cancer classification. It has also used a deep neural network (DNN) to solve simple linear classifiers. Based on DeepGene’s [1] experiences and reports, we examined three deep learning models for cancer classification, with the CNN model showing the best results with 66.64% test accuracy. Figure 4 shows the structure and components of this model. Figure 11 and Figure 12 show the LSTM and CNN+LSTM models, respectively.

**Figure 4.**
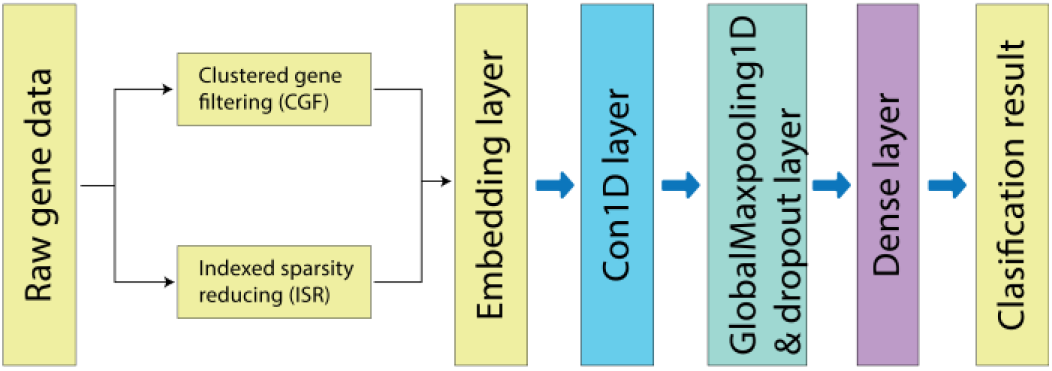
our proposed CNN model diagram.

We used the ISR pre-processing and embedding layer to eliminate data sparsity. To identify the genes involved in the classification work, we used CGF pre-processing and CNN. We used two regularization methods to solve the overfitting problem, Dropout, and Max-norm. Since several fully connected layers are susceptible to overfitting in the CNN network, we used the global pooling layer to solve the overfitting problem in the CNN model. To solve the problem of using simple linear classifiers, we used a dense layer.

The CNN model steps are as follows: First, we convert raw data to binary (zero and one). Second, ISR and CGF [1] accept this converted data as input. Then ISR output concatenates to the tail of CGF output, which is used as input for the embedding layer. CNN layer extract features after receiving the output of the embedding layer and throws the data to global max pooling. At the end of the layer, a dense layer is responsible for classifying the data received from global max pooling.

We used the database introduced by Yuan et al.[1], in all our experiments, a re-formulated subset of The Cancer Genome Atlas (TCGA). In the following, we will describe each of the mentioned components.

### 4.2 Pre-processing

There are two pre-processing steps, CGF and ISR, descript by Yuan et al. [1] CGF aims to create a subset of genes with a higher frequency of mutations, thus eliminating genes that do not give us valuable information. For this reason, the First step is to calculate the sum of all the mutations for each gene and sort them descending. CGF keeps the index of all genes in this step. In the second step, genes are grouped by similarity Jaccard coefficient, if a group has more than five-member, then ISR keeps the five genes that have the most mutation in the group, and if a group has less than five members, CGF deletes that group. All selected genes in this part create a matrix that is the output of CGF.

The goal of the ISR is to eliminate the vast number of zeros in the data, which reduces the complexity of the gene data. For each sample, ISR selects all genes that have a mutation in that sample and keeps their index. If the number of these genes is greater or equal to 800, then ISR selects the top 800 genes with the highest mutation, but if the number of genes is less than 800, then ISR adds zero to the end of the vector. This process is applied to all samples. Then the results create a matrix that is the output of ISR.

To apply these two pre-processors to the data, we must first convert the data to binary (zeros and ones). One means that the gene has mutated, and zero means that the gene did not mutate. Then we apply each of these two methods separately to the database. We attach the output matrix from the ISR to the end of the output matrix from the CGF and use this new matrix as input to the CNN, LSTM, and CNN and LSTM combination. Due to the individual performance of CGF and ISR, our data has substantially zero values, which affects the performance of our models. We suggested an embedding layer to solve this problem, as explained below.

### 4.3 Embedding Layer

Our input data given to the model has zero, one, and several index numbers obtained in the ISR step. Despite the use of ISR and CGF pre-processing, the number of zeros in our data is high, and it causes problems for our models. To solve this problem in our three models, we first give our data to the embedding layer. Keras Library created this embedding layer. This layer can also solve the sparsity problem in the matrix. This layer takes positive integer numbers as input and creates a dense vector of constant length for each number. This vector has weights that the Keras generated randomly. Keras update embedding vectors during the training process [67, 68]. Suppose we have a vector [0, 1, 2, 3, 4]. When we give it to the embed layer and want to create vectors of length two, the result will be in Table 1.

**Table 1.** Map integer index is to the embedding vectors.

| Index | Embedding |
| --- | --- |
| 0 | [2.1, 3.2] |
| 1 | [0.5, 3.4] |
| 2 | [1.2, 1.4] |
| 3 | [0.7, 1.7] |
| 4 | [3.0, 7.1] |

The Table 1 shows that the embedding layer connects each of the numbers inside the original vector to a vector of constant length two. Zero map to [2.1, 3.2], one map to [0.5, 3.4], etc. Now we have new vector like this, [[2.1, 3.2], [0.5, 3.4], [1.2, 1.4], [0.7, 1.7], [3.0, 7.1]].

#### 4.3.1 Embedding Layer mask-zero parameter

It is prevalent in sequence data processing that some samples do not have the same length, some are shorter than others, and some are longer than others. To make the length of the data the same, we add zero to the end of the data to make the length of the samples equal that occurs called padding. The ISR method, which we mentioned in the pre-processing section, does the same thing to equalize the length of the samples. After unifying the length of the samples using padding, the model must declare and ignore that part of the data is padding. We do this by setting the TRUE value for the mask zero parameters of the Embedding layer created by the Keras library that will improve the performance of the LSTM network in this article.

### 4.4 Dropout and Global max pooling

A convolutional network comprises convolution, pooling, and fully connected layers Section. 3.2. In this way, the convolution and pooling layers are located at the beginning of the network. All these layers play the role of feature extraction for our model. In the end, there are fully connected layers, which end with a softmax or logistic regression layer classification is placed [69]. Because fully connected layers are prone to overfitting, they can cause problems throughout the network.

Hinton et al. [70] Suggest regularization methods such as the dropout layer used to prevent overfitting. This layer randomly deactivates several neurons in each epoch, only done during the training phase. We can also replace the fully connected layers traditionally used in CNN network architecture with the global max-pooling layer. In this way, we create a feature map for each of the relevant categories for classification in the last layer. We take the maximum of each feature map and pass the current vector directly to softmax [71, 72]. If the global max pooling input is as follows: 0, 1, 2, 3, 4, 5, 1, 2, the output will be 5 [73].

### 4.5 Data collection

To fairly compare the results of our models and the model proposed by Yuan et al. [1], we use their suggested database, which is a re-formulated subset of the Cancer Genome Atlas (TCGA) [74] database called TCGA-DeepGene. In this dataset, Yuan et al. have collected genes containing somatic point mutations in each of the 12 types of cancer. Gene mutations are stored in this database in binary form; zero means genes not mutated, and one means gene mutated. The database has 22834 genes and 3122 samples, the dimensions of the database are 22834 by 3122. Each of the dataset columns is labeled 1 to 12 because it has 12 different types of cancer. Table 2 shows details of samples and mutations in each type of cancer [1].

**Table 2.** Samples and mutation statistic of the TCGA-DeepGene [1] dataset.

|  | Sample number | Missense mutation | Nonsense mutation | Nonstop mutation | RNA | Silent | Splice Site | Translation start site | Total mutation |
| --- | --- | --- | --- | --- | --- | --- | --- | --- | --- |
| ACC | 91 | 6741 | 501 | 15 | 368 | 2534 | 344 | 42 | 10545 |
| BLCA | 130 | 24067 | 2142 | 46 | 0 | 9662 | 528 | 55 | 36500 |
| BRCA | 992 | 55063 | 4841 | 133 | 3998 | 17901 | 1424 | 0 | 83360 |
| CESC | 194 | 26606 | 2716 | 84 | 5595 | 9765 | 527 | 0 | 45293 |
| HNSC | 279 | 31416 | 2545 | 44 | 0 | 12149 | 776 | 0 | 46930 |
| KIRP | 171 | 8910 | 499 | 17 | 394 | 3411 | 524 | 0 | 13755 |
| LGG | 284 | 5341 | 378 | 7 | 102 | 2074 | 294 | 0 | 8196 |
| LUAD | 230 | 44800 | 3477 | 46 | 0 | 15594 | 1377 | 99 | 65393 |
| PAAD | 146 | 21067 | 1496 | 19 | 859 | 7936 | 1005 | 111 | 32493 |
| PRAD | 261 | 9628 | 563 | 15 | 652 | 3750 | 513 | 55 | 15176 |
| STAD | 288 | 82265 | 4200 | 92 | 48 | 3334 | 1868 | 227 | 122044 |
| UCS | 56 | 3070 | 187 | 2 | 234 | 1114 | 171 | 0 | 4778 |
| Total | 3122 | 318974 | 23545 | 520 | 12250 | 119234 | 9351 | 598 | 484463 |

## 5. Experiments and results

In this section, we detail the steps taken to implement our models and evaluate the performance of each model to select the best of them.

### 5.1 Prototypes implementation

We have implemented our deep neural network models in Python. The Python libraries, Pandas and scikit-learn, are used to read the dataset, build the two pre-process and data splitting. We used the Tensorflow platform and the Keras API to build our models, CNN presented in Figure 4, LSTM shown in Figure 11, and CNN+LSTM shown in Figure 12. Keras has introduced the Keras-tuner library to adjust the hyper-parameters mentioned in section 5.3, which we also used to achieve our goal. Due to the lack of data, and the small number of samples, we need to measure the model performance accurately. Also, different samples in the training set can affect the model training process and its accuracy rate on the test data, so the model should be trained and evaluated with different parts of the data. 10-fold cross-validation ensures that any observation of the original data set has a chance to appear in the training and validation set. 10-fold cross-validation shows the average accuracy rate of the model after using all folds. Also, we can select the model that has the highest accuracy in 10-fold cross-validation and use it on test data to get the highest accuracy rate. In this way, we have the model mean accuracy and the model which trained well on the training dataset. [75–78]. After setting the hyper-parameters, we use the technique of 10-fold cross-validation, which we have created with the help of the scikit-learn library to measure the model’s performance on the train and test dataset. In all of our experiments, we used Sparse Categorical Cross entropy as a loss function provided by Keras API.

### 5.2 Data splitting and Hyper-parameters

After CGF and ISR pre-processing, we divided our data into train, test, and validation datasets. We create a train and test set using the train-test-split method from the sklearn library. In this distribution, the train-test-split method separates 10% of the total data for the test dataset and 90% for the training dataset. We use 10% of the training dataset for the validation test in our training phase.

For the 10-fold cross-validation, we split the data into train and test with the rate we mentioned above, then trained the model on the train set and used the fold-out part for the validation set; after the training step, we used the test dataset to test the model. We want to know what combination of the hyper-parameter values listed below can increase each of our three models’ performance. The names and values of the hyper-parameters we want to test are listed below.

output-dim for the Embedding layer between numbers [1, 2, 4, 8, 16, 32, 64, 128, 256, 512, 1024], dropout rate, for the layer dropout between numbers [0.5, 0.6, 0.7, 0.8, 0.9], max-value, for max-norm between numbers [1, 2, 3, 4], learning rate between numbers [0.1, 0.01, 0.001, 0.0001], number of units for LSTM layer between numbers [1, 5, 10, 15, 20, 25, 30, 35, 40, 45, 50], the number of filters for CNN between numbers [2, 4, 8, 16, 32, 64, 128, 512], the Keras library has been used to implement all layers.

### 5.3 Hyper-parameter tuning

To set our hyper-parameters, we used the train set. For this purpose, we use the random search method offered on the Keras-tuner library. We set an important parameter for this library: max-trials to twenty, executions-per-trials to two, and validation split to 10%. Due to the imbalance of the dataset shown in Table 2, we use the stratified k-fold method in the sklearn library. This method causes each set to contain approximately the same percentage of each class sample as the complete set.

### 5.4 Create and tun CNN experiment

In this section, we describe how to tune and test the CNN model. In the first three steps, the CNN model used the training dataset.

In section 1, we said there are four main steps to building and adjusting all three models. We will set all the hyper-parameters mentioned in Section 5.3 in these four steps for the CNN model. **Step one**, we will adjust the learning rate, the number of filters, and the Embedding layer output-dim using Keras-tuner. Table. 3 shows the top ten of the best settings. We choose the best Values for learning rate = 0.001, the number of filters = 512, and embedding layer output-dim = 512. In Section 4.1, we declared that we had used an embodied layer to improve the performance of our models.

**Table 3.** ten of best values for hyper-parameters. We chose to tune in the first step of creating the CNN model.

| Filters | Output dim | Learning rate | Validation accuracy |
| --- | --- | --- | --- |
| 512 | 512 | 0.001 | 0.6601 |
| 512 | 64 | 0.01 | 0.6245 |
| 256 | 1024 | 0.0001 | 0.5907 |
| 128 | 1024 | 0.01 | 0.5658 |
| 16 | 32 | 0.01 | 0.5658 |
| 16 | 128 | 0.0001 | 0.411 |
| 512 | 4 | 0.001 | 0.4092 |
| 256 | 4 | 0.01 | 0.40 |
| 16 | 1024 | 0.001 | 0.3967 |
| 64 | 4 | 0.001 | 0.3825 |

To investigate the effect of the embedding layer on the performance of the CNN model, we compare the model’s performance with and without embedding layer with hyper-parameters in Step one. For this reason, we remove the embedding layer from the model and retrain the CNN model, then check the validation accuracy of the CNN model with and without the embedding layer. By comparing Figures 5a and 6a together, we can see the difference between the validation accuracy of 60% for the CNN model using the Embedding layer and 35% validation accuracy for the CNN model without the embedding layer. Also, Figures 5b and 6b show the difference in loss between the CNN model that uses the embedding layer validation loss of 1.5 and the CNN model without embedding layer validation loss of 500. These significant differences between validation accuracy and validation loss encouraged us to use the embedding layer in other stages to manufacture CNN.

**Figure 5.**
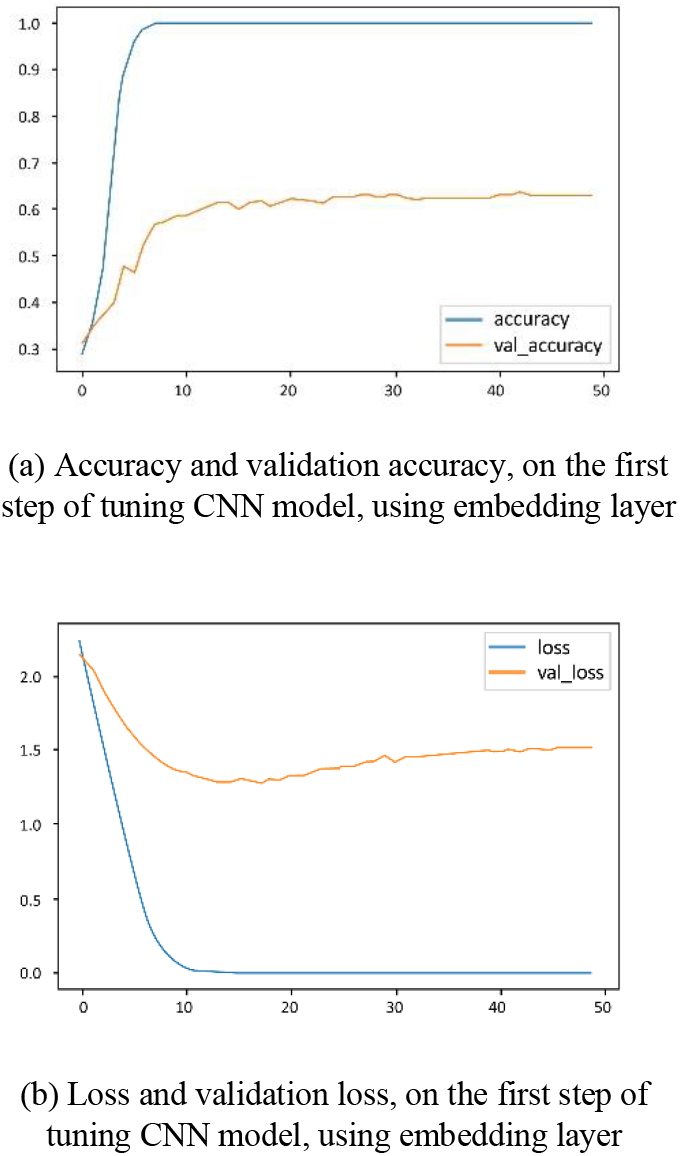
Accuracy and Loss in First Step of Create CNN Model with Embedding Layer.

**Figure 6.**
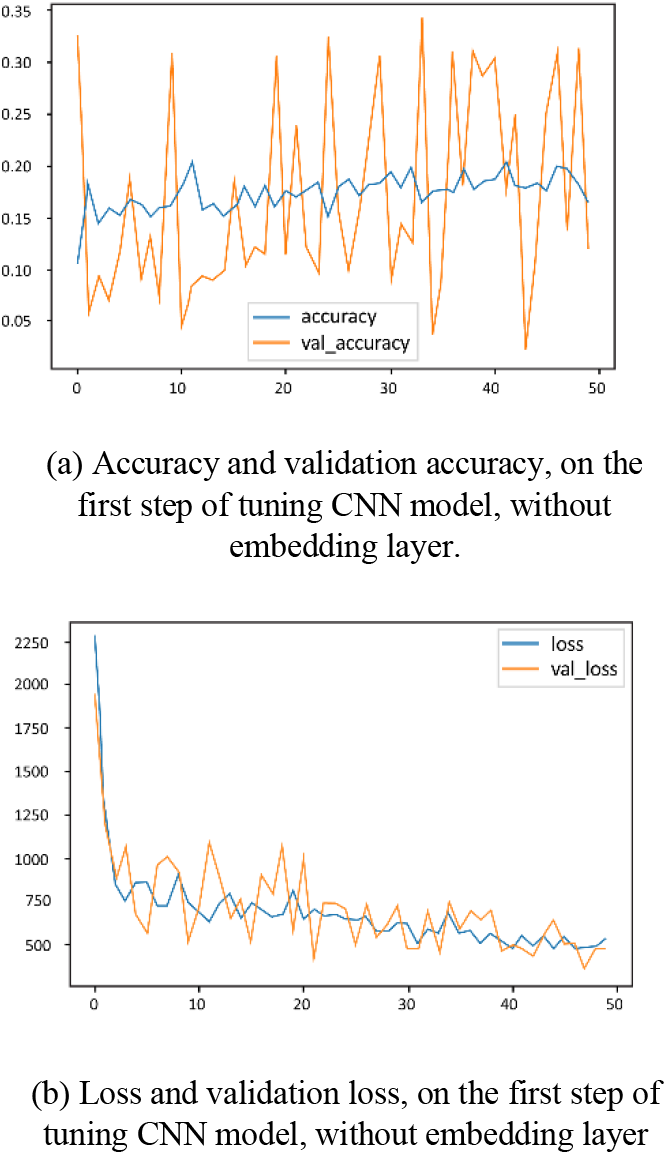
Accuracy and Loss in First Step of Create CNN Model without Embedding Layer

Figure 5b shows a big difference in train loss and validation loss after epoch 15, indicating overfitting. **In step Two**, to solve the overfitting problem, we add two dropout layers, dropout-one between the embedding layer and Conv-1D, dropout-two between global max pooling and dense layer. Then use Keras-tuner to get the best values for dropout-rate and max-norm. We do not change the hyper-parameters that we find in the first step of the tuning. Table 4 shows the ten best values for these parameters. As can be seen, the best validation accuracy is 0.7135 with max-norm = 2.0, dropout-one = 0, and dropout-two = 0.5. Because dropout-one is zero, we remove this layer in our CNN structure. By comparing Figure 5b and Figure 7b, we can see the model performance is improving. According to Tables 3 and 4, this improvement from 66.01% to 71.35% after setting new hyper-parameters in step two. In Figure 5b, the overfitting starts from epoch 15, but in Figure 7b, overfitting starts from epoch 20.

**Table 4.** ten of best values for hyper-parameters for solve CNN model overfitting problem. Second Step of Tuning CNN.

| Dropout One | Dropout Two | Max-norm | Validation accuracy |
| --- | --- | --- | --- |
| 0 | 0.5 | 2.0 | 0.7135 |
| 0 | 0.6 | 2.0 | 0.6939 |
| 0 | 0.8 | 1.0 | 0.6921 |
| 0 | 0.5 | 4.0 | 0.6921 |
| 0 | 0.7 | 3.0 | 0.6921 |
| 0 | 0.6 | 4.0 | 0.6921 |
| 0.5 | 0.5 | 1.0 | 0.6868 |
| 0 | 0.8 | 2.0 | 0.6850 |
| 0 | 0.7 | 4.0 | 0.6832 |
| 0.5 | 0.8 | 3.0 | 0.6797 |

**Figure 7.**
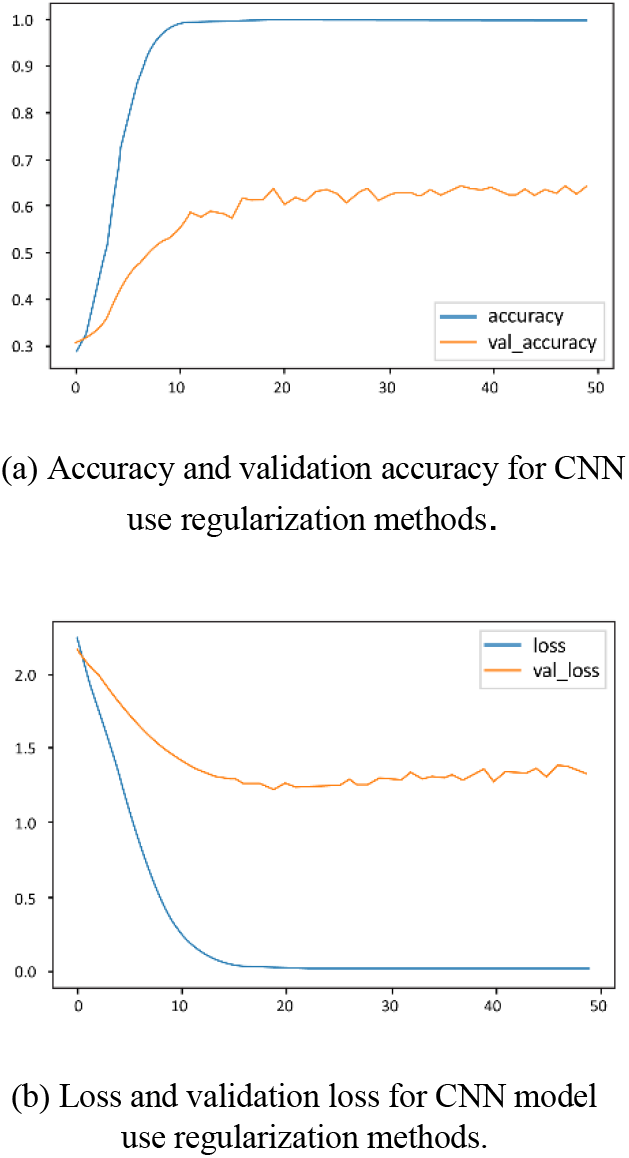
Loss and validation loss for CNN model use regularization methods.

After tuning steps one and two, we have the best hyper-parameters for each layer of CNN structure. We are also sure that the embedding layer increases the performance of the CNN model. **In step three**, with the help of the stratified k-fold function, we implement the 10-fold cross-validation method. For this reason, we used our training dataset to split it into ten folds, nine folds used for the training step, and one fold used for the validation step. We will monitor the validation accuracy in each fold and store the weights that give us the most validation accuracy in the training phase. **Step four**, after the 10-fold cross-validation, we will load these weights on the CNN model structure and test them on the test dataset. We report the best test accuracy as the final result. Table 5 shows these results.

**Table 5.**
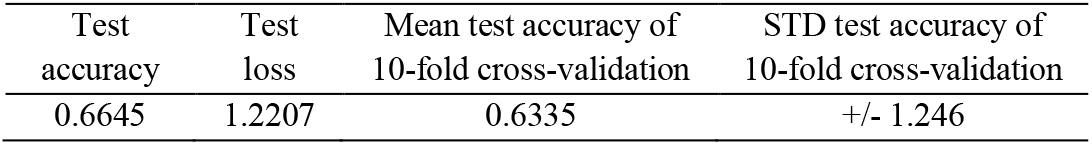
result of stratified 10-fold cross validation and best model of all folds on test dataset for CNN model

| Test accuracy | Test loss | Mean test accuracy of 10-fold cross-validation | STD test accuracy of 10-fold cross-validation |
| --- | --- | --- | --- |
| 0.6645 | 1.2207 | 0.6335 | +/- 1.246 |

### 5.5 Evaluation Metrics

In order to better observe the performance of our chosen model (CNN) on the test data, we calculated the criteria precision, recall, and F1-score. Table 6 shows the classification report, and Figure 8 is a confusion matrix.

**Table 6.**
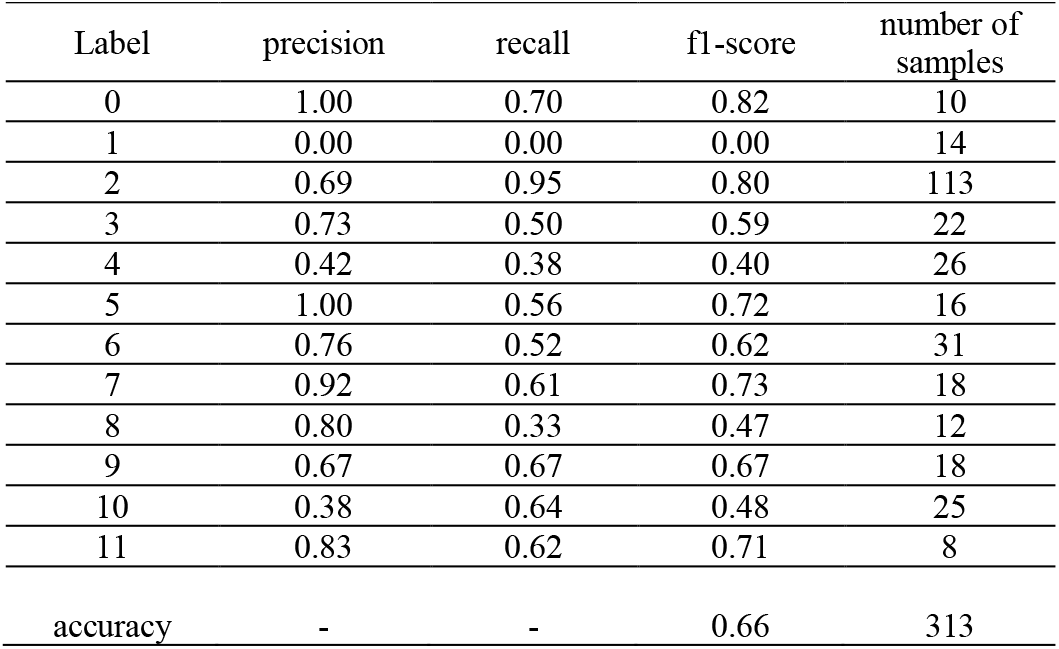
Classification report for proposed model (CNN)

**Figure 8.**
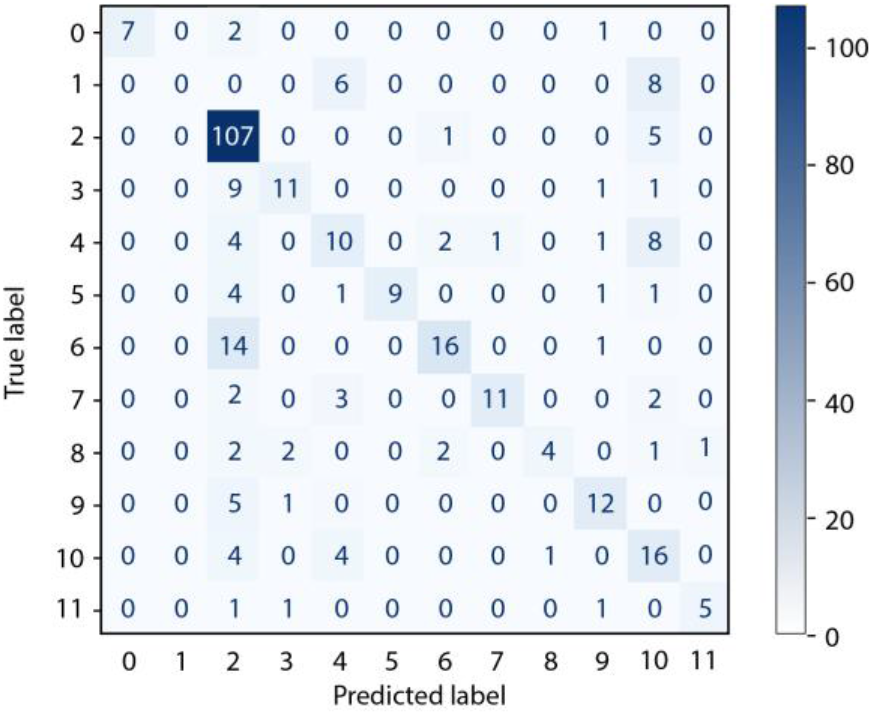
Confusion matrix visualization for our proposed model (CNN)

According to Table 6, our labels were numbered from 0 to 11, according to Table 2, ACC, BLCA, BRCA, CESC, HNSC, KIRP, LGG, LUAD, PAAD, PRAD, PRAD, STAD, UCS, respectively. For calculating the precision, recall, and f1-score, first, we need to calculate true positive (TP), true negative (TN), false positive (FP), and false-negative (FN), then calculate precision, recall, and f1-score for each label. The formula (4) [79, 80] show how to calculate these criteria for one label. Table 6 shows that labels zero and two have the highest value of f1-score, and label one has the lowest value of f1-score. Label number one is mainly confused with labels number 4 and 10 due to the small number of samples and similarity of features. Label number two has the largest sample size in the total data, making it better to identify the model. Despite the small number of samples, Label number zero has more unique features than label one, which has increased its f1-score. According to Figure 8, in the classification task, the model made mistakes in predicting the labels. Labels are often mistaken for labels 2, 9, and 10. Ten labels were misidentified by two, six labels were misidentified by 9, and 6 labels were misidentified by 10. A large number of features compared to the number of samples has caused the similarity between the features to be high. Due to the small number of samples, model in training and validation stages have problems finding unique features for each class and classification.

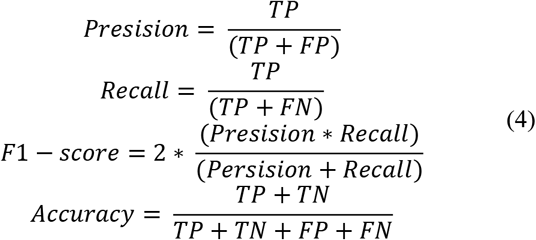

### 5.6 Create and tune LSTM experiment

In this section, we will talk about setting up and building an LSTM model. In the first three steps, the LSTM model used the training dataset.

**In the first step**, we adjust the learning rate, the number of cells, and the value of the output parameter for the embedding layer using a Keras-tuner. Table 7 shows the ten best values for setting these hyper-parameters, for which we select the values learning rate = 0.01, the number of cells = 64, and embedding layer output-dim = 1024.

**Table 7.** Ten of best values for hyper-parameter for solve LSTM model overfitting problem. Second step of LSTM tuning model

| Dropout | Max-norm | Validation accuracy |
| --- | --- | --- |
| 0.6 | 2.0 | 0.4644 |
| 0.5 | 4.0 | 0.4626 |
| 0.5 | 3.0 | 0.4572 |
| 0 | 4.0 | 0.4551 |
| 0.5 | 2.0 | 0.4537 |
| 0 | 3.0 | 0.4537 |
| 0.6 | 3.0 | 0.4519 |
| 0.7 | 2.0 | 0.4519 |
| 0.8 | 4.0 | 0.4483 |
| 0 | 1.0 | 0.4483 |

To show the effect of the embedding layer on the performance of the LSTM model, we remove this layer from the model. We keep the learning rate value and the number of cell hyper-parameters obtained above and trained the model. Then we compare the value of validation accuracy obtained from the model with the embedding layer and the model without embedding layer. LSTM model without the embedding layer reduces the validation accuracy from 44.48% to 37.90%. Due to this reduction in the validation accuracy, we will use the embedding layer in other stages of setting up and manufacturing LSTM.

Figure 9 shows the accuracy and loss of the LSTM model in the first step of tuning. As shown in Figure 9b, there is a big gap between train loss and validation loss, which is a sign of an overfitting problem. Our validation loss in 50 epochs is going to 6. **In the second step**, to solve this problem of tuning LSTM, we use the hyper-parameters in the first step, then adjust the dropout parameters and max-norm using a Keras-tuner. Table 8 shows ten of the best values obtained for these hyper-parameters. We select the values 0.6 for dropout and 2 for max-norm. Figure 10 shows the accuracy and loss in the second step of tuning LSTM. In compression with Figure 9, there is a significant difference between validation loss in Figure 9b and Figure 10b. Our validation loss in Figure 9b goes up to 6. However, in Figure 10b, our validation loss goes up to 4 in the same period. Also, comparing Figure 9a and Figure 10a, validation accuracy increased slightly from 44.48% to 46.44%. This result shows us that the hyper-parameters that we tuned in the second step help the model reduce the validation loss and increase validation accuracy.

**Table 8.** ten of the best settings for hyper-parameters in the first step to build the LSTM model.

| Cell | Output dim | Learning rate | Validation accuracy |
| --- | --- | --- | --- |
| 64 | 1024 | 0.01 | 0.4448 |
| 32 | 32 | 0.01 | 0.427 |
| 32 | 64 | 0.01 | 0.4234 |
| 32 | 512 | 0.001 | 0.4163 |
| 32 | 4 | 0.001 | 0.395 |
| 32 | 256 | 0.0001 | 0.3843 |
| 16 | 1024 | 0.0001 | 0.3807 |
| 2 | 32 | 0.01 | 0.3807 |
| 4 | 1024 | 0.001 | 0.3754 |
| 16 | 4 | 0.01 | 0.3736 |

**Figure 9.**
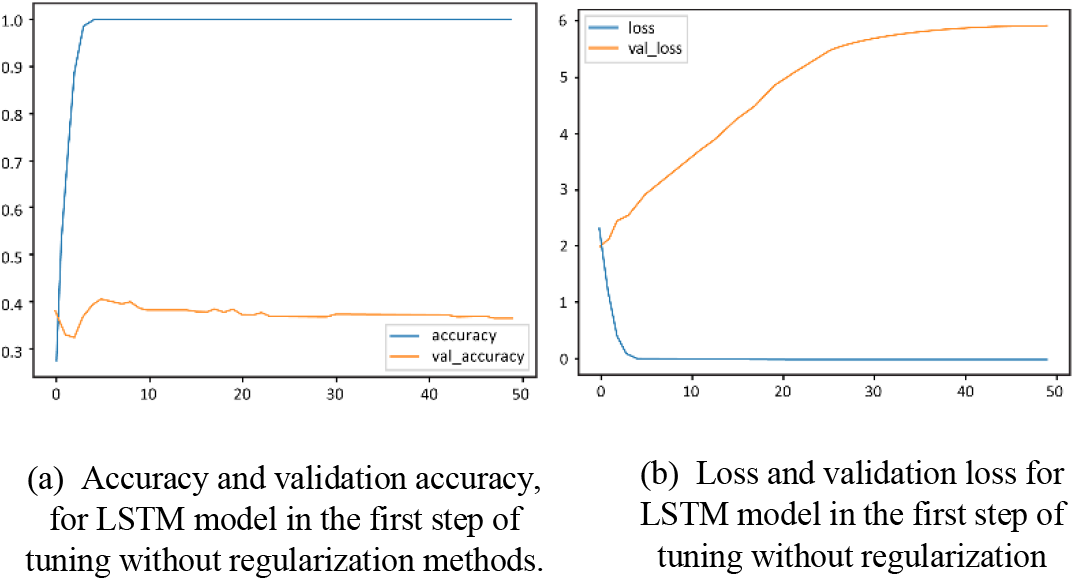
train and validation accuracy and loss for LSTM model in tt second step of tuning with regularization methods

**Figure 10.**
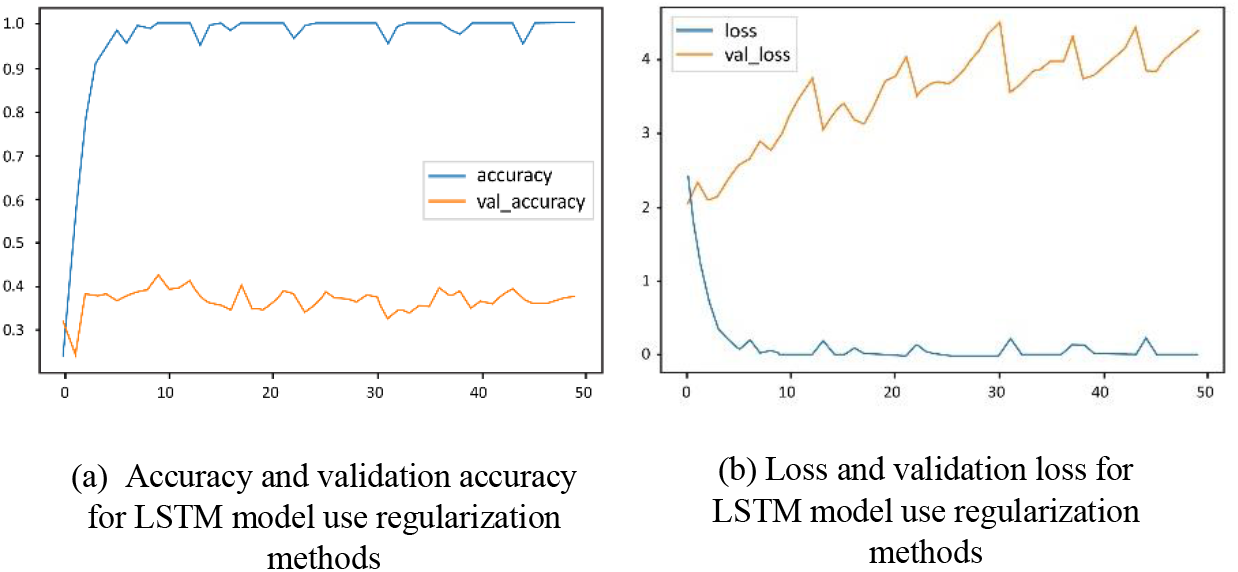
Train and validation loss and accuracy for LSTM model in the first step of tuning without regularization methods

We obtained the best values for the parameters mentioned above in the previous two steps. We will implement the 10-fold cross-validation method using the stratified k-fold left out for the validation test **in the third step**. We store weights that give us the best validation accuracy in each fold. We load these weights on our LSTM model and test it using a test dataset **in step four**. Table 9 shows the best test accuracy and Figure 11 shows the structure of our LSTM model.

**Table 9.**
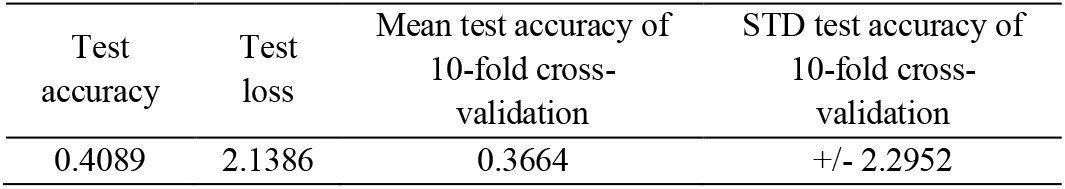
result of stratified 10-fold cross validation and best model of all folds on test dataset for LSTM model

### 5.7 Create and tune CNN + LSTM

We use the best values of the hyper-parameters specified in sections 5.4 and 5.5 for the LSTM and CNN models combination. Using the stratified k-fold function, we implement the 10-fold cross-validation method and train the CNN+LSTM model on the training dataset. We store the best weights that have the most validation accuracy in each fold. After completing the 10-fold cross-validation, we load the weights on the model and test it with the test dataset. Figure 12 shows the structure of the CNN+LSTM model. Table 10 shows the test accuracy of our three models. Also, Figure 13 compares our model test accuracy with DeepGene deep learning and machine learning methods. Figure 13 shows that the CNN model has the highest test accuracy compared with the fully connected (DeepGeen) created by [1]. Furthermore, CNN have positive impact to LSTM performance, simple LSTM test accuracy = 40.89% but CNN + LSTM = 41.2%. Machine learning techniques implemented by Yuan et al [1] support vector machine (SVM) have the highest test accuracy compared with K-Nearest Neighbor (KNN) and Naive Bayes (NB).

**Table 10.** Accuracy of all models on the test dataset.

| Model | Accuracy |
| --- | --- |
| CNN | 0.6645 |
| LSTM | 0.4089 |
| CNN + LSTM | 0.4120 |

**Figure 11.**
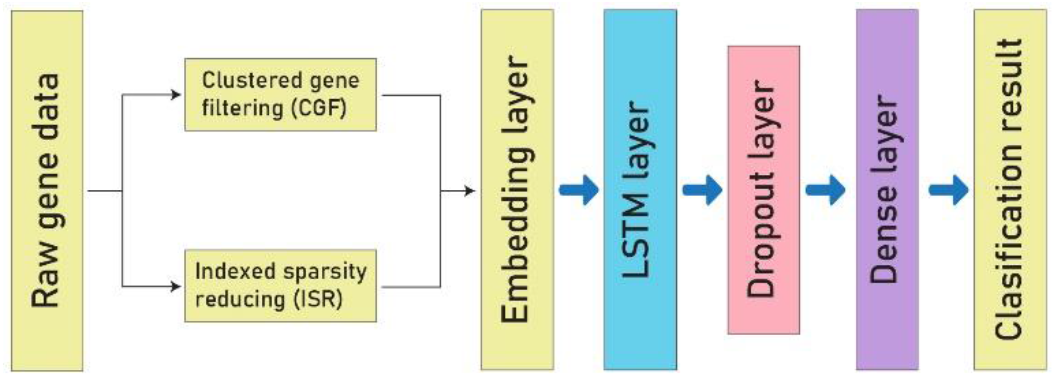
Final structure of LSTM model.

**Figure 12.**
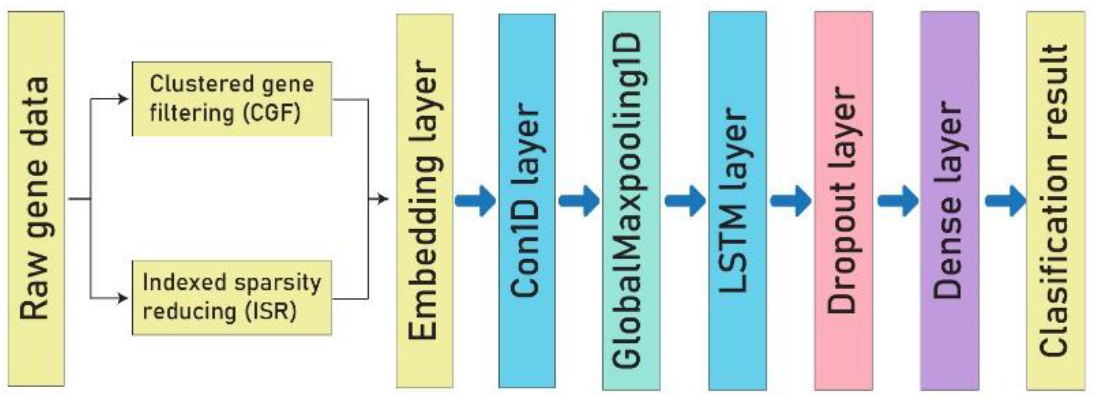
structure of CNN + LSTM model.

**Figure 13.**
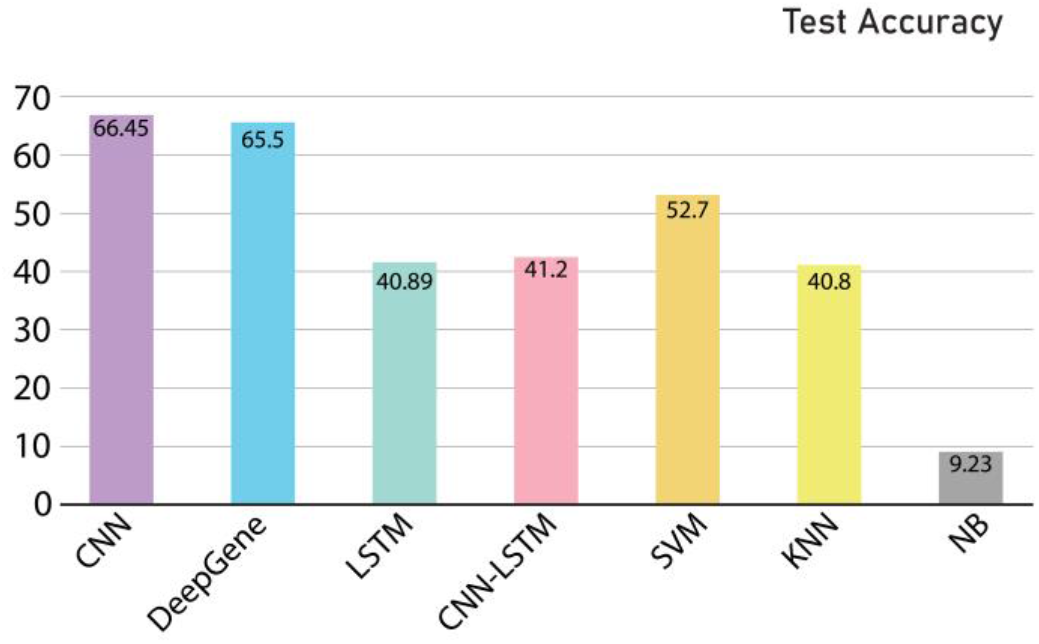
Comparison of CNN, LSTM, CNN+LSTM test accuracy with DeepGene deep learning and machine learning methods SVM, KNN, and NB.

### 5.8 DeepGene implementation

To compare the performance of the approach provided and Yuan et al.[1], we tried to execute the code they write by MATLAB programming language in the Github repository. Due to the lack of an important folder and its contents, we could not execute the code and reproduce the results they declared.

That is why we implemented their model with Python programming language in the same way they described in their article. Table 11 shows the best test accuracy obtained from stratified 10-fold cross-validation. We were unable to reproduce the values using the information provided by Yuan et al. [1]

**Table 11.** Result of DeepGene Implementation.

| State | Max accuracy | Mean accuracy of 10-fold cross-validation | STD test accuracy of 10-fold cross-validation |
| --- | --- | --- | --- |
| Train | 0.3285 | 0.3191 | +/- 0.316 |
| Test | 0.3258 | 0.3175 | +/- 0.287 |

## 6. Discussion

SMCC methods promise to find more accurate and faster ways for cancer type/subtype identification. Using machine learning techniques in SMCC can create new and effective strategies for diagnosing cancer. The simple linear classifiers used in machine learning, the sparsity of data, and many genes in a DNA sequence are some of the problems that machine learning systems have with somatic point data. Addressing these challenges can make cancer diagnosis more accurate, faster, and cheaper. It can also pave the way for new treatments.

In this study, based on the results reported by DeepGene [1] and used the pre-processing and datasets introduced by them, we test three deep learning models for cancer classification. Of these three models, CNN shows the best result. CNN is a deep learning model that has made significant advances in medical images to diagnose cancer and cancerous tumors. CNN also has a high ability to maintain valuable features and remove unnecessary elements. Figure 4 shows the structure of our proposed model.

Our experiments show that the embedding layer significantly reduced sparsity data and improved our proposed model’s performance. The dropout and max-norm layers also impacted reducing the effects of overfitting in the CNN model. The CNN model and the combination of CNN and LSTM have had better results than the LSTM model, which shows that CNN has an excellent performance in selecting features. In combination with other models, CNN can increase its performance in this area of research.

We found that to improve the performance of deep learning models in this field of research. We need more robust pre-processing methods to identify essential features and thus increase the quality of data input to the model. Combined with other deep learning or machine learning models, the CNN model can also increase its performance by identifying influential features in classification work. We need models that are resilient to overfitting due to insufficient data.

## 7. Conclusion

The diversity and prevalence of cancers and the dramatic growth of those with the disease have made cancer one of the leading causes of human death. Genomics based technologies have provided a great opportunities for researchers to study genomic variants, including somatic point mutations. Developing techniques that can use genomic s information to detect cancer types early, accurately, and quickly has become an urgent need. Machine learning methods have recently used somatic mutations to identify cancer subtypes and subtype-specific biomarkers [1, 35, 38, 81–83]. This article tested three deep learning models and the pre-processors introduced in DeepGene [1] for cancer classification. CNN showed the highest accuracy on the test dataset of the three models tested. We found that adding an embedding layer to the CNN model increased its performance. Also, by comparing the results of LSTM models and combining CNN and LSTM, we realized that the CNN model combined with other models could improve their performance.

We intend to work on pre-processing methods that identify effective genes and thus increase the quality of data input to the model for future work.

## Data Availability Statement

The dataset analyzed during the current study are available in the DeepGene repository, https://github.com/yuanyc06/deepgene.

## Author contributions

HAR and PP designed the study; PP, MF, MR designed the models. PP, HAR, MF wrote the paper. HAR, MF, MR, and PP edited the manuscript. PP carried out all the analyses, including the statistical analyses, model developments, comparision, etc. PP generated all figures and all tables. All authors have read and approved the final version of the paper.

## Acknowledgments

Analysis was made possible with computational resources provided by the BioMedical Machine Learning high performance computing Server with funding from the Australian Government and the UNSW SYDNEY.

